# Abundance drives broad patterns of generalisation in plant-hummingbird pollination networks

**DOI:** 10.1101/339762

**Authors:** Benno I. Simmons, Jeferson Vizentin-Bugoni, Pietro K. Maruyama, Peter A. Cotton, Oscar H. Marín-Gómez, Carlos Lara, Liliana Rosero-Lasprilla, María A. Maglianesi, Raul Ortiz-Pulido, Márcia A. Rocca, Licléia C. Rodrigues, Boris Tinoco, Marcelo F. Vasconcelos, Marlies Sazima, Ana M. Martín González, Jesper Sonne, Carsten Rahbek, Lynn V. Dicks, Bo Dalsgaard, William J. Sutherland

## Abstract

Abundant pollinators are often more generalised than rare pollinators. This could be because abundance drives generalisation: neutral effects suggest that more abundant species will be more generalised simply because they have more chance encounters with potential interaction partners. On the other hand, generalisation could drive abundance, as generalised species could have a competitive advantage over specialists, being able to exploit a wider range of resources and gain a more balanced nutrient intake. Determining the direction of the abundance-generalisation relationship is therefore a ‘chicken-and-egg’ dilemma. Here we determine the direction of the relationship between abundance and generalisation in plant-hummingbird pollination networks sampled from a variety of locations across the Americas. For the first time we resolve the direction of the abundance-generalisation relationship using independent data on animal abundance. We find evidence that hummingbird pollinators are generalised because they are abundant, and little evidence that hummingbirds are abundant because they are generalised. Additionally, a null model analysis suggests this pattern is due to neutral processes: most patterns of species-level abundance and generalisation were well explained by a null model that assumed interaction neutrality. These results suggest that neutral processes play a key role in driving broad patterns of generalisation in animal pollinators across large spatial scales.

**Declarations:** Funding – BIS is supported by the Natural Environment Research Council as part of the Cambridge Earth System Science NERC DTP [NE/L002507/1]. JVB was funded by CERL - Engineer Research and Development Center. PKM was funded by the São Paulo Research Foundation (FAPESP grant #2015/21457-4). PAC was funded by the David Lack studentship from the British Ornithologists’ Union and Wolfson College, University of Oxford. CL was funded by the ESDEPED-UAT grant. MAM acknowledges the Consejo Nacional para Investigaciones Científicas y Tecnológicas (Costa Rica), German Academic Exchange Service and the research funding program ‘LOEWE-Landes-Offensive zur Entwicklung Wissenschaftlichö konomischer Exzellenz’ of Hesse’s Ministry of Higher Education, Research, and the Arts (Germany). ROP was funded by CONACyT (project 258364). MAR was supported by the State of São Paulo Research Foundation (FAPESP) within the BIOTA/FAPESP, The Biodiversity Institute Program (www.biota.org.br) and the ‘Parcelas Permanentes’ project, as well as by Coordenação de Pessoal de Nível Superior (CAPES), Fundo de Apoio ao Ensino e à Pesquisa (FAEP)/Funcamp/Unicamp and The Nature Conservancy (TNC) of Brazil. LCR was supported by CNPq and Capes. MS was funded by CNPq (grant #302781/2016-1). AMMG is supported through a Marie Skłodowska-Curie Individual Fellowship (H2020-MSCA-IF-2016-704409). LVD was supported by the Natural Environment Research Council (grants NE/K015419/1 and NE/N014472/1). AMMG, JS, CR and BD thank the Danish National Research Foundation for its support of the Center for Macroecology, Evolution and Climate (grant no. DNRF96). WJS is funded by Arcadia.

## Introduction

Pollination and other mutualistic associations are crucial for the functioning and maintenance of ecological communities (Heithaus 1974, Rech et al. 2016, Ollerton 2017, Ratto et al. 2018). A common phenomenon in mutualistic communities is that more abundant species tend to have more generalised interaction niches, interacting with a greater number of partners than rare species (Dupont et al. 2003, Vázquez and Aizen 2003, Olesen et al. 2008). Thus, abundant species are often ‘ecological’ generalists (Ollerton et al. 2007). However, the direction of the relationship between abundance and generalisation has been described as a ‘chicken-and-egg’ dilemma as there are valid *a priori* explanations for both directions (Fort et al. 2016, Dormann et al. 2017). For example, high abundance could lead to high generalisation simply due to neutral effects: more abundant species have a higher likelihood of encountering a greater number of potential interaction partners than rarer species (Vázquez et al. 2007, 2009, Poisot et al. 2015). Additionally, pollinators have been observed to increase their generalisation when at high densities: in a given area, higher species abundance leads to greater conspecific competition for the available resources, resulting in increased generalization as predicted by optimal foraging theory (Fontaine et al. 2008, Tinoco et al. 2017). Conversely, high generalisation could lead to high abundance. For example, the wider diet breadth of generalist individuals could be advantageous in communities with high levels of variability or species turnover where flexibility is beneficial (Waser et al. 1996, CaraDonna et al. 2017). Such ‘portfolio effects’ allow a mutualist to receive a more stable benefit over time despite having partners with asynchronous dynamics or different performance trade-offs (Batstone et al. 2018). Generalisation can also provide a better nutrient balance (Tasei and Aupinel 2008, Behmer 2009, Vaudo et al. 2015), improve species’ pathogen resistance (Alaux et al. 2010, Di Pasquale et al. 2013) and afford functional redundancy that buffers against partner extinction (Biesmeijer et al. 2006). In a recent review by Batstone et al. (2018), the authors argue that generalisation in mutualisms can have a selective advantage over specialisation for many reasons. These include sampling effects, where generalisation increases the likelihood that a given mutualist will sample the most beneficial partner (for example, Albrecht et al. 2012), and complementarity, where a given mutualist benefits from having diverse partners that occupy different niches, but provide the same rewards via different mechanisms. Therefore, generalization may confer advantages to specific pollinators, resulting in higher abundances.

Here we evaluate the direction of the abundance-generalisation relationship in plant-hummingbird pollination networks and use a null model to assess the extent to which observed patterns of species-level generalisation can be explained by neutral effects. We focus on hummingbird species, rather than plants, as plants may have non-hummingbird mutualistic partners not included in our data that could result in misleading estimates of generalisation (Dalsgaard et al. 2008). Plant-hummingbird interactions are a particularly interesting model system to answer these questions as they involve species spanning the entire specialisation-generalisation spectrum (Bleiweiss 1998, Martín González et al. 2015, Dalsgaard et al. 2018) and recent studies suggest that abundance has little influence on network structure compared to morphological trait matching (Maruyama et al. 2014, Vizentin-Bugoni et al. 2014, 2016, Weinstein and Graham 2017, though see Bergamo et al. 2017 and Dalsgaard et al. 2018). Understanding the processes governing specialisation and generalisation can also contribute to explaining the high diversity of sympatric hummingbird species as niche partitioning can play a role in species coexistence, thus it may also add to the explanation why there are so many species in the tropics (Dalsgaard et al. 2011). Additionally, pollination by vertebrates is important, especially in the tropics (Bawa 1990, Vizentin-Bugoni et al. 2018), and is on average responsible for 63% of fruit or seed production in vertebrate-pollinated plants (Ratto et al. 2018). Therefore, understanding the abundance-generalisation relationship in vertebrate pollinators such as hummingbirds has important implications for understanding the processes maintaining tropical plant and vertebrate communities. While previous attempts to resolve the abundance-generalisation chicken-and-egg dilemma have used species’ interaction frequency as a proxy for animal abundance (Fort et al. 2016), which can lead to biased conclusions (Vizentin-Bugoni et al. 2014), here we use independent animal abundance estimates. This is an important advance because all 35 pollination and seed dispersal networks analysed by Fort et al (2016) used estimates of animal abundance based on the interaction network data, and the authors had direct measures of plant abundance for only 29% of networks. By their own admission, “These animal abundance data are arguably limited, as they are not independent from the interactions; but these are the best data available to evaluate our question.” Conversely, ours is the first study where we have estimates of plant and animal abundance independent from the interaction observations for the majority of networks. This study also represents a significant methodological advance for resolving the chicken-and-egg dilemma. While Fort et al (2016) classified species’ abundance and generalisation using either strict thresholds or parametric fuzzy logic methods which assume linearity, our approach makes no such assumptions about the distributions of the data and uses the data’s full continuous range without the use of thresholds. We find evidence of a unidirectional relationship with hummingbird abundance driving hummingbird generalisation. Importantly, a null model assuming neutrality of interactions closely matched most empirical results. This suggests that neutral effects have an important role in structuring broad patterns of species-level generalisation, even in a system such as plant-hummingbird pollination networks where phenotypical matching has a strong influence on the occurrence of pairwise interactions among species.

## Material and Methods

### Dataset

We assembled a database of plant-hummingbird pollination networks with complementary information on hummingbird and plant abundance. In total we gathered 19 quantitative networks, where link weights represent the number of observed hummingbird visits to plants. In total, the database contained 103 hummingbird species and 403 plant species. For each of the 19 networks, hummingbird abundances were quantified as the mean number of individuals per species either recorded along transect counts within the sampling plots or caught using mist nets (Appendix 1). For four networks where species were not recorded within the sampling plots during transect counts or mist netting, we used frequency of occurrence (the proportion of days of fieldwork in which a given species was recorded) as a proxy for relative abundances, as both measures are strongly correlated and frequency of occurrence is still independent from the network data (Vizentin-Bugoni et al. 2014). To test whether these four networks affected our results, we repeated all analyses excluding these data (Appendix 2). Plant abundances were quantified along transect counts or inside plots within the study areas and summarized as the number of flowers per species recorded over the sampling period. Species abundances and interactions were quantified several times (typically, monthly) over at least a complete annual cycle in each community. Further details of each network are given in Appendix 1.

### Measures of generalisation

We calculated the level of generalisation of all hummingbird species in all networks. To assess the sensitivity of our results to the choice of generalisation metric, we measured generalization in three ways. First, species degree, which is simply the number of plant species a given hummingbird species interacts with. Second, normalised degree, which is equal to a species’ degree divided by the total number of possible partners. Third, a generalisation index *g*, based on a widely used species-level measure of specialization (*d*′) that quantifies the extent to which a species deviates from a random sampling of its available interaction partners (Blüthgen et al. 2006). We calculated *d*′ using independent abundance data. To ensure that higher values of *d*′ corresponded to higher levels of generalisation, we calculated the standardised generalisation index *g*, defined as 1-*d*′/*d*′max where *d*′max is the maximum possible value of *d*′ (Fort et al. 2016). *d*′ and *d*′max were calculated using the ‘dfun’ function in the ‘bipartite’ R package (Dormann et al. 2009).

### General approach

First, we tested whether there was a relationship between hummingbirds’ abundance and their level of generalisation using three linear mixed effects models, one for each generalisation metric. The generalisation metric was the response variable, with log(abundance) as a fixed effect and species and network identity as random effects. A Poisson distribution was used for the model with degree as the response variable, a binomial distribution was used for the model with normalised degree as the response variable (with weights equal to the maximum degree of each species) and a Gaussian distribution was used for the model with *g* as the response variable. Mixed effects models were fitted using the ‘lme4’ R package (Bates et al. 2015) and the significance of fixed effects was calculated using Wald χ^2^ tests available in the ‘Anova’ function of the ‘car’ R package (Fox and Weisberg 2002). We calculated both the marginal R^2^_(G)LMM(m)_, which represents the variance explained by fixed effects, and the conditional R^2^_(G)LMM(c)_, which represents the variance explained by both fixed and random effects (Nakagawa and Schielzeth 2013, Emer et al. 2016, Kaiser-Bunbury et al. 2017, Bartoń 2018).

Having established that there is a relationship between abundance and generalisation, we used the approach of Fort *et al.* (2016) to determine whether abundance drives generalisation or generalisation drives abundance. This approach uses formal logic, specifically material implication, to derive expectations for broad species-level patterns of abundance and generalisation in ecological communities. To explain the approach, it is useful to consider a simple example. Consider the proposition, *P*, “if it is a dodo, it is extinct”. *P* is made up of two statements: (i) “it is a dodo” and (ii) “it is extinct”. Given that each of these statements can either be true or false, we can derive four possible outcomes, as shown in Table 1. Outcome A is a dodo that is extinct. Outcome B is a non-dodo that is not extinct, such as the hummingbird species *Amazilia versicolor*. Outcome C is a non-dodo that is extinct, such as the dinosaur species *Tyrannosaurus rex*. Finally, outcome D is a dodo that is not extinct. We can only refute the proposition “if it is a dodo, it is extinct” when we observe outcome D to be true; that is, if we observe a living dodo. Conversely, observing an extinct dodo, an extant *Amazilia versicolor* individual, or an extinct *T. Rex* specimen are all consistent with *P*.

**Table 1:** Truth table listing all possible outcomes for the propositions “if it is a dodo, it is <u>extinct” and “if it is abundant, it is generalist”. ‘T’</u> is ‘True’ and ‘F’ is ‘False’.

| Outcome | Dodo/Abundant | Extinct/Generalist |
| --- | --- | --- |
| A | T | T |
| B | F | F |
| C | F | T |
| D | T | F |

There are four possible outcomes when applying this to the abundance-generalisation chicken-and-egg dilemma: abundant generalists, rare generalists, abundant specialists and rare specialists (Table 1). We can therefore derive two hypotheses:

1. If abundance implies generalisation, there should be no species which are abundant and specialist (outcome D: living dodos); we would only expect to observe abundant generalists (outcome A: extinct dodos), rare specialists (outcome B: a living *Amazilia versicolor*) and rare generalists (outcome C: extinct *T. Rex*).
2. If generalisation implies abundance, there should be no generalist species that are rare; we would only expect to observe rare specialists, abundant specialists and abundant generalists.

Therefore, by calculating the proportion of hummingbird species in each of the four abundance-generalisation categories (rare specialists, abundant specialists, rare generalists and abundant generalists), it is possible to test these two hypotheses and determine whether the relationship between hummingbird abundance and generalisation is unidirectional (Fort et al. 2016). Particularly it is important to look at the proportion of rare generalists and abundant specialists: if hypothesis 1 is correct, there should be few abundant specialists; if hypothesis 2 is correct, there should be few rare generalists.

### Abundance and generalisation classification

To calculate the proportion of hummingbird species in each abundance-generalisation category, we developed a novel methodology to classify each species in a community as either rare or abundant and as either specialist or generalist. As mentioned above, this improves on Fort et al’s (2016) methodology by making no assumptions about the distributions of the data and by using the data’s full continuous range without the use of thresholds. For each network, we first rescaled the abundance and generalisation values of all hummingbird species to range between 0 and 1 according to (*x* – *x*_min_)/(*x*_max_ – *x*_min_), where *x*min and *x*max are the minimum and maximum values of abundance or generalisation (Aizen et al. 2012). These values represent the probability with which a species would be classified as abundant or generalist. Next, we sampled a random value from a uniform distribution between 0 and 1. If a species’ rescaled abundance or generalisation was greater than or equal to this value, it was classified as abundant or generalist, respectively. If it was less than this value, it was classified as rare or specialist, respectively. Therefore, a species with a rescaled abundance of 0.2 would have a 20% probability of being classified as abundant in a given iteration. Similarly, a species with a rescaled abundance of 0.8 would have an 80% probability of being classified as abundant. This was repeated 1000 times. The mean proportion of species in each of the four abundance-generalisation categories was then calculated. This was repeated for each of the three generalisation metrics.

### Null model analysis

To assess the extent to which our results could be explained purely by neutral effects, we used a null model to generate 1000 randomised versions of each empirical network. The null model assumed interaction neutrality by assigning interactions according to a probability matrix, **A**, where element *a_ij_* was the relative abundance of hummingbird species *i* multiplied by the relative abundance of plant species *j* (Vázquez et al. 2007, Maruyama et al. 2014, Vizentin-Bugoni et al. 2014, 2016). Therefore, the model assumes that two species with high abundance have a greater likelihood of interacting than two species with low abundance. The model constrained the number of links and ensured that each species had at least one interaction (Vázquez et al. 2007). We used independent plant and hummingbird abundance data to create the null networks, rather than relying on species marginal totals as a proxy for abundance. For each of the 1000 null versions of each of the 19 empirical networks, we repeated the permutational analysis described above (‘Abundance and generalisation classification’) to calculate the mean proportion of species in each of the four abundance-generalisation categories predicted by the neutral model. We then compared these proportions based on neutrality to the empirical proportions: if the empirical proportions were within the 95% confidence intervals of the null model proportions then there were no significant differences between the null model and the observed values.

## Results

We confirmed the positive relationship between abundance and generalisation in our dataset, finding a significant correlation between abundance and generalisation for degree (Wald test: χ^2^ = 216.44; df = 1; *P* = < 0.001; R^2^_GLMM(m)_ = 0.55; R^2^_GLMM(c)_ = 0.79), normalised degree (Wald test: χ^2^ = 232.1; df = 1; *P* = < 0.001; R^2^_GLMM(m)_ = 0.26; R^2^_GLMM(c)_ = 0.37) and the generalisation index *g* (Wald test: χ^2^ = 10.7; df = 1; *P* = 0.001; R^2^_LMM(m)_ = 0.06; R^2^_LMM(c)_ = 0.44).

Only a small proportion of species were abundant and specialist for all three generalisation metrics (Figure 1). Conversely, the proportion of species that were rare and generalist was consistently larger, particularly for the *g* generalisation metric. These differences were significant: the proportion of species that were rare and generalist was significantly higher than the proportion which were abundant and specialist for degree (t = 2.92, df = 18, *p* = 0.009), normalised degree (t = 2.91, df = 18, *p* = 0.009) and *g* (t = 10.34, df = 18, *p* = < 0.001) (Figure 1). Overall, these findings support hypothesis 1, that abundance drives generalisation, and do not support hypothesis 2, that generalisation drives abundance.

**Figure 1:**
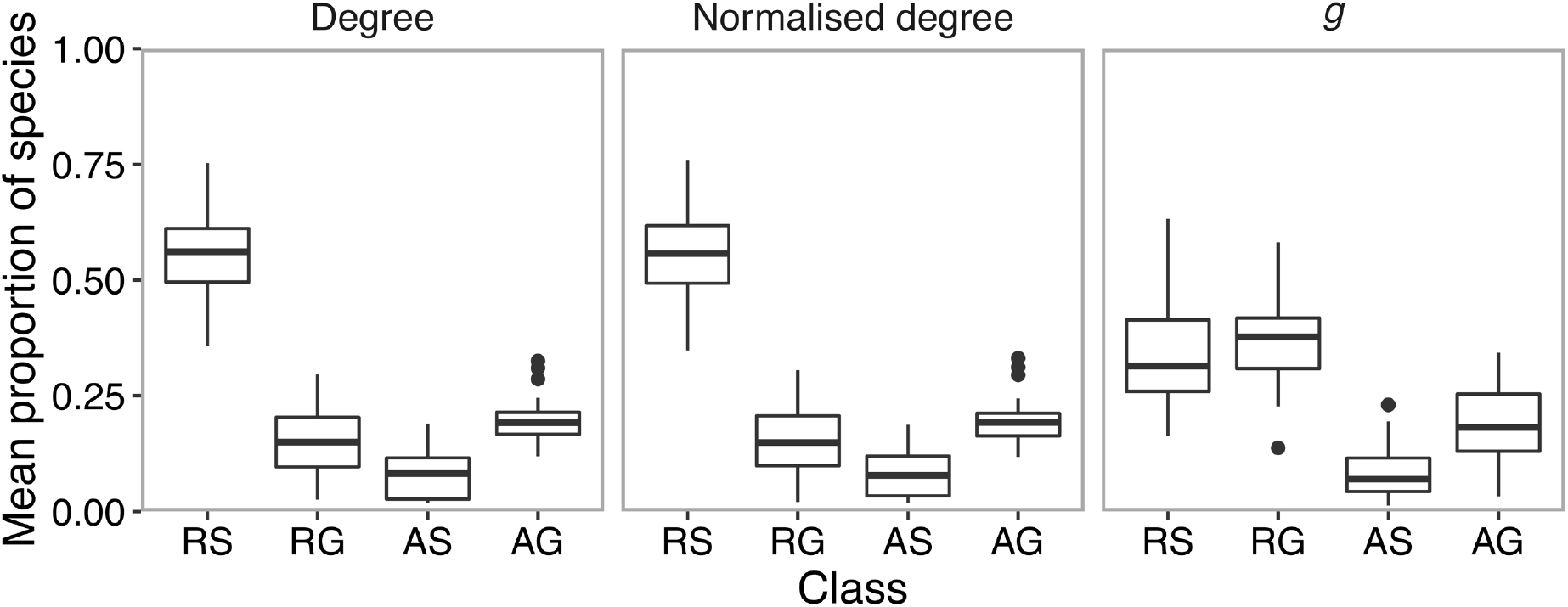
The mean proportion of hummingbird species classified as rare specialists (‘RS’), rare generalists (‘RG’), abundant specialists (‘AS’) and abundant generalists (‘AG’) across all networks, for three generalisation metrics: degree, normalised degree and *g*. The bold centre line in each box is the median; the lower and upper hinges are the first and third quartiles, respectively. The lower whisker indicates the smallest value no less than 1.5 times the inter-quartile range; the upper whisker indicates the largest value no greater than 1.5 times the inter-quartile range. Data outside the whiskers are outlying points plotted as solid black circles.

The proportion of species in each of the four abundance-generalisation categories predicted by the neutrality null model closely matched the empirical proportions, particularly for degree and normalised degree where there were no significant differences between observed and predicted proportions for the majority of networks (68–84% of networks; Figure 2). For *g*, the model correctly predicted the proportion of rare specialists and generalists for 79% of networks, but performed less well in predicting the proportion of abundant specialists and generalists, with predictions matching observed values for only 47% of networks (Figure 2).

**Figure 2:**
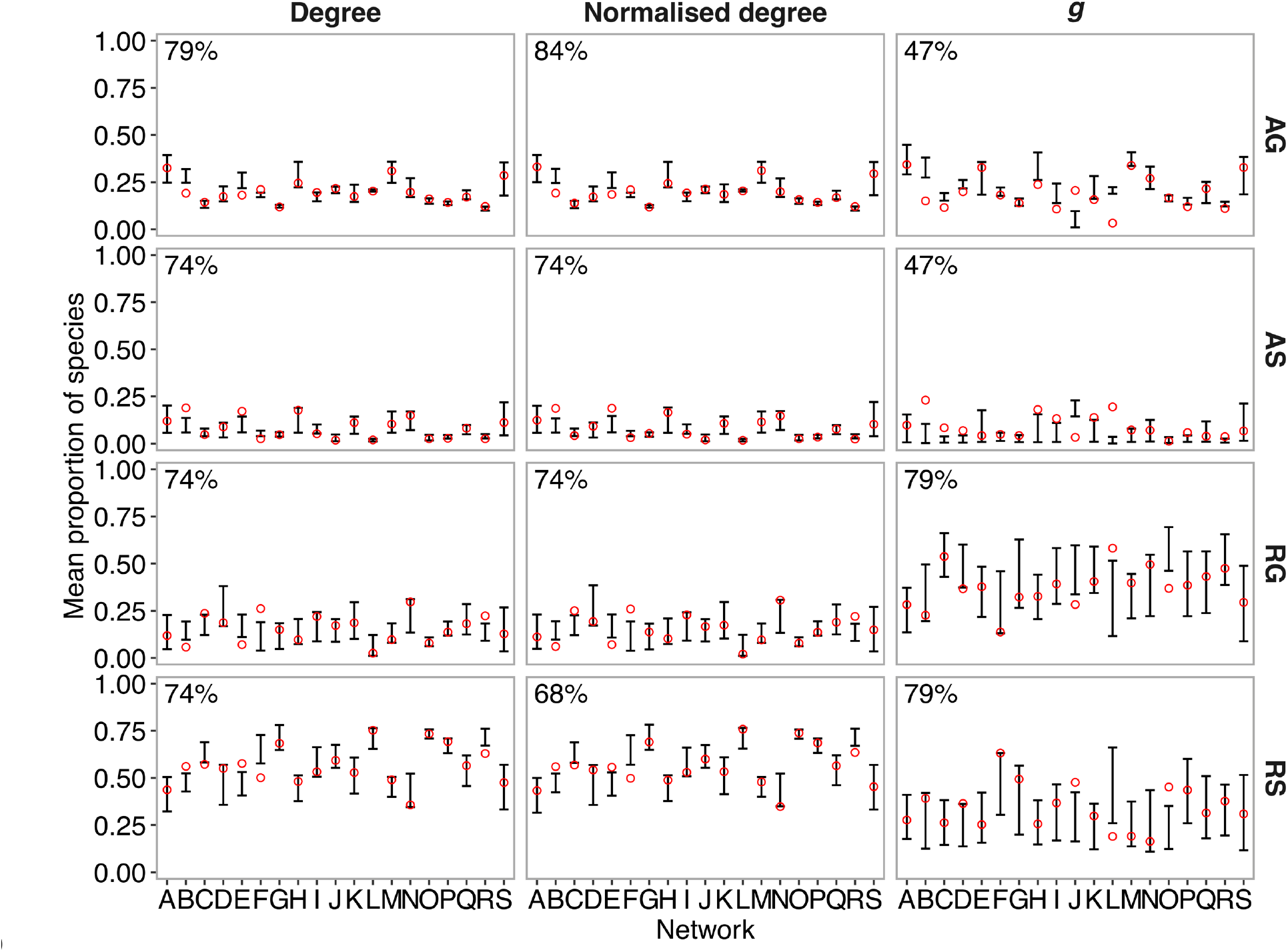
Comparisons between empirical networks (A-S) and null model networks in the proportions of species in each of the abundance-generalisation categories ‘RS’ (rare specialists), ‘RG’ (rare generalists), ‘AS’ (abundant specialists) and ‘AG’ (abundant generalists). Error bars represent the 95% confidence intervals of the mean proportion of hummingbird species in each abundance-generalisation category as predicted by 1000 null networks. Red circles show the empirically observed mean proportion of hummingbird species in each category. If the red circle is within the error bars, there were no significant differences between the observed proportions and the neutrality null model proportions. Percentages in the top left of each panel give the proportion of networks where empirical proportions were not significantly different from the null model proportions. Results are shown for each network (A-S) and for each generalisation metric (Degree, Normalised degree, *g*).

All results were qualitatively the same and conclusions identical after the exclusion of the four networks where we used frequency of occurrence (the proportion of days of fieldwork in which a given species was recorded) as a proxy for relative abundances (Appendix 2).

## Discussion

Our analysis of numerous plant-hummingbird communities sampled widely across the Americas support the hypothesis that abundance drives species-level generalisation, and provide little evidence that generalisation drives species abundance. These results can be discussed in the context of *sufficient* and *necessary* conditions from formal logic. If we say that *P* is a *necessary* condition for *Q*, then in the absence of *P* there is also an absence of *Q*. For example, sitting the exam is a *necessary* condition for getting an A grade. If a student does not sit the exam, they will not get an A grade. Similarly, if a student is awarded an A grade, they must have sat the exam. However, if *P* is a *sufficient* condition for *Q*, then if we have *P*, *Q* must follow. For example, obtaining full marks on every exam question is a *sufficient* condition for getting an A grade in the exam. Therefore, if a student gets full marks on every question, they will get an A grade. However, getting full marks on every question is not a *necessary* condition for getting an A grade: it is possible to get an A without achieving full marks on all questions. Similarly, sitting the exam is not a *sufficient* condition for getting an A grade: it is possible to sit the exam and not get an A. Our results suggest abundance is a *sufficient* condition for generalisation as, if a species is abundant, it tends to also be a generalist. However, it is not a *necessary* condition as species can be generalist without being abundant. Conversely, our results suggest generalisation is a *necessary* condition for abundance as, if a species is a specialist, it tends to be rare. However, it is not a *sufficient* condition for abundance as, if a species is a generalist, this does not mean it is abundant. Therefore, our results agree with those of Fort *et al.* (2016) using pollination and seed dispersal networks, suggesting that abundance driving generalisation may be a general phenomenon that can be observed in mutualistic systems.

In all ecological studies it is worth asking whether sampling effort may impact the results. This is also the case for studies of species interaction networks, as sampling effects can influence the observed network structure (Fründ et al. 2016, Jordano 2016, Vizentin-Bugoni et al. 2016, Dalsgaard et al. 2017). Sampling is likely to result in missed detections of interactions for rare species, resulting in an underestimation of how generalised rare species are (Blüthgen 2010, Dorado et al. 2011). For this reason, Dormann et al. (2017) described sampling rare species with high generalisation as “impossible”. This means that our results are unlikely to be a function of sampling effects, as the proportion of rare generalist species we observe is likely less than the true proportion: under theoretical perfect sampling, we would likely observe a larger proportion of species which are rare generalists, reinforcing our results (Dorado et al. 2011). Furthermore, sampling effects are likely to overestimate the proportion of species that are rare specialists as, even when rare species are observed, they are unlikely to be observed on all the plants they visit. This suggests that sampling effects will cause the generalisation level of rare species to be underestimated, and that consequently some species classified as rare specialists may actually be rare generalists (Blüthgen 2010, Dorado et al. 2011). Sampling effects are therefore not likely to impact our conclusions, because with perfect sampling we would expect the proportion of rare generalists to increase and the proportion of rare specialists to decrease, further increasing support for hypothesis 1 (many rare generalists, few abundant specialists) and refuting hypothesis 2 (few rare generalists, many abundant specialists). Additionally, we would not expect sampling artefacts to explain the low proportion of species which were abundant specialists because sampling effects tend to come from missing links for rare species rather than abundant species (Blüthgen 2010, Dorado et al. 2011, Fort et al. 2016).

A frequent interpretation of the abundance-generalisation relationship is that abundant species are more generalised due to neutral effects; that is, they are more likely to encounter a greater number of interaction partners than less abundant species by chance alone (Vázquez et al. 2007). Our null model analysis supports this interpretation, particularly for degree and normalised degree: we found that the numbers of rare specialists, abundant specialists, rare generalists and abundant generalists were well predicted for the majority of networks by a null model that assumed interactions were formed entirely from neutral processes. This finding complements other recent studies of plant-hummingbird pollination networks showing the importance of morphological trait matching in predicting pairwise interactions at the network level (Maruyama et al. 2014, Vizentin-Bugoni et al. 2014, 2016, Weinstein and Graham 2017), while here we show that abundance predicts broad patterns of generalisation at the species level. Among Antillean hummingbirds, it was recently shown that local environmental conditions and floral richness, not hummingbirds’ morphological traits, determined species level nectar-feeding specialization (Dalsgaard et al. 2018). Combined with our findings, this might suggest a hierarchy of mechanisms structuring plant-hummingbird interactions, and more broadly whole pollination networks (Junker et al. 2013, Bartomeus et al. 2016, Vizentin-Bugoni et al. 2018): neutrality and local conditions govern broad patterns of generalisation, such as the number of plant partners, while morphological matching operates at a lower level to determine the identity of these plant partners. For the generalisation index *g*, the null model performed less well, predicting the proportion of abundant specialists and abundant generalists correctly in only 47% of networks. For the remaining 53% of networks, the model generally over predicted the number of abundant generalists and under predicted the number of abundant specialists. This may be due the nature of the *g* index itself: by accounting for the abundance of plants, *g* does not necessarily correlate with species degree (number of plant partners). For example, a hummingbird which visits one abundant plant could receive a higher value of *g* than a hummingbird that visits three rare plants. This means the null model may overestimate the number of abundant generalists and underestimate the number of abundant specialists as, in the model, an abundant hummingbird will have a higher probability of interacting with all plants, while in the empirical network it may be able to gain sufficient resources by only interacting with the most abundant plants.

Taken together, our study confirms that abundance is a sufficient, but not necessary, condition for generalisation in plant-hummingbird pollination networks; it is the first study to test this hypothesis in animals using independent data on species abundance encompassing a wide array of communities. Remarkably, our result corroborates the findings of Fort et al. (2016), giving further support that this may be a general phenomenon in mutualistic systems. Further research should investigate whether the relationships found here hold for other types of ecological systems. We also find evidence that neutral effects are good predictors of coarse species-level patterns of generalisation, even in a system in which interactions are widely recognized to be constrained by species traits. This might suggest a hierarchy of mechanisms structuring plant-hummingbird interactions, with neutral effects operating at a ‘high level’ to determine coarse patterns of generalisation, such as the number of partners, while niche-based processes act at a lower level to determine the identity of these partners.

## Data accessibility

Data will be deposited in Data Dryad before we submit a revised version of the manuscript prior to acceptance.

